## Supporting Information for "Structure and mechanism of the RNA dependent RNase Cas13a from *Rhodobacter capsulatus*"

### Structural and mechanistic insights into Cas13a from *Rhodobacter capsulatus*

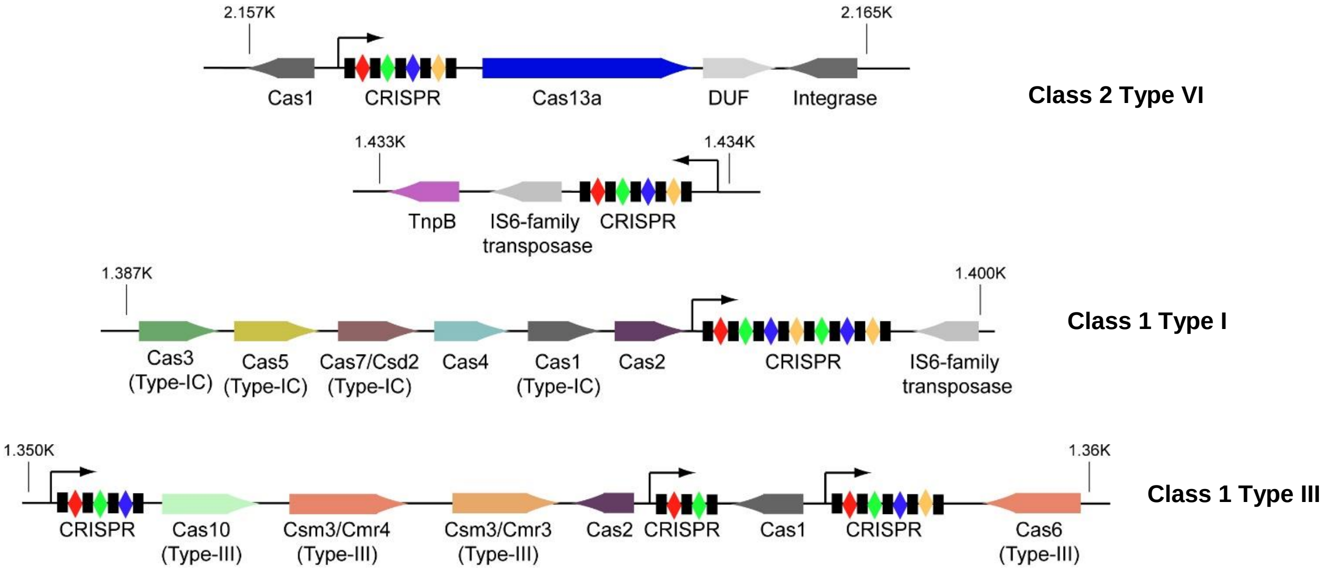


**Supporting Figure S1.** Genomic location and structure of CRISPR-Cas sequences identified in the reference genome of *Rhodobacter capsulatus SB1003* (NCBI accession number NC_014034.1) using the CRISPR-finder (1) and BLAST search (2).

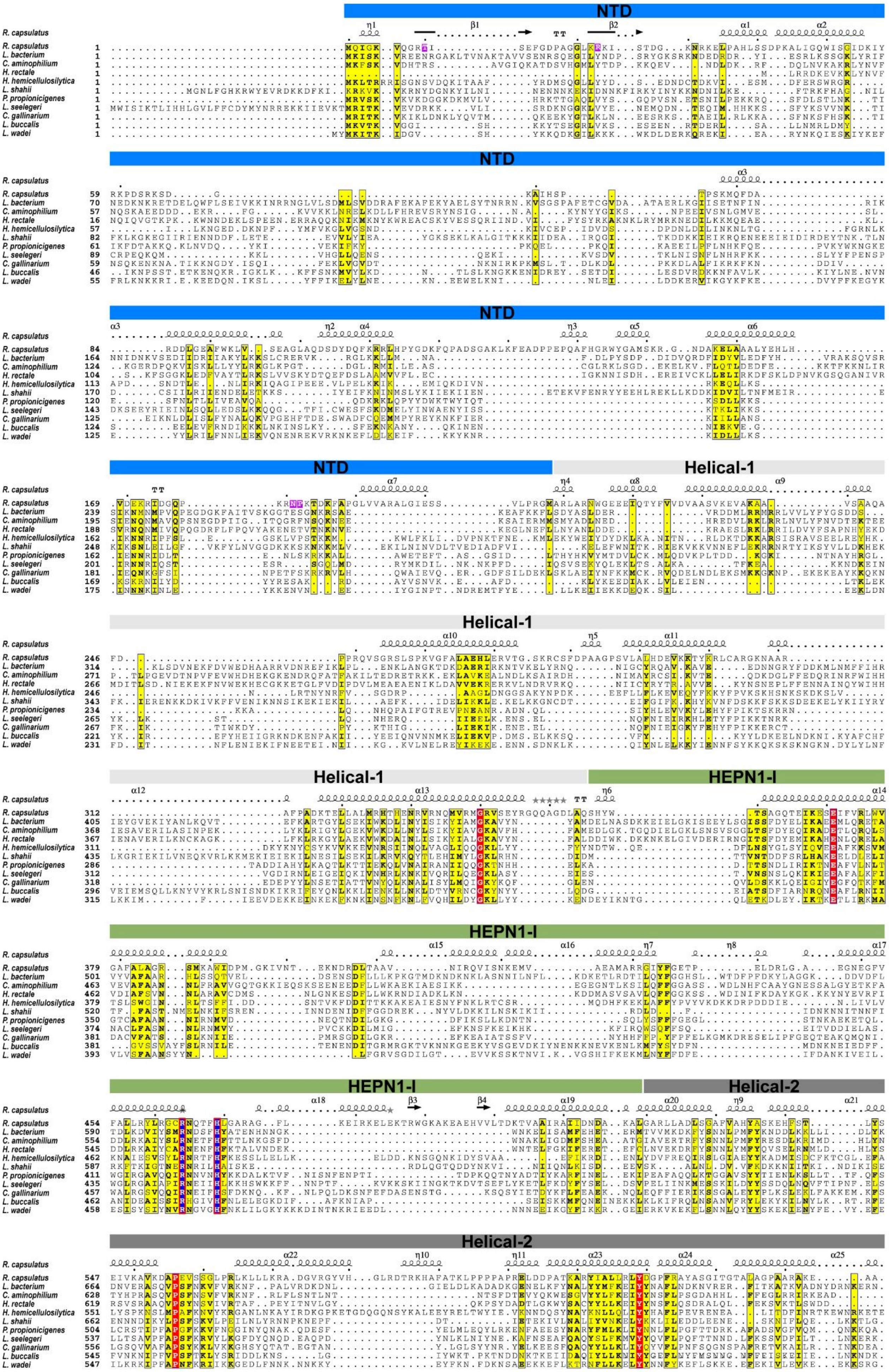
NTD
Helical-1
HEPN1-I
Helical-2
R. capsulatus
L. bacterium
C. aminophilium
H. rectale
H. hemicellulosilytica
L. shahii
P. propionicigenes
L. seegeri
C. gallinarium
L. buccalis
L. wadei

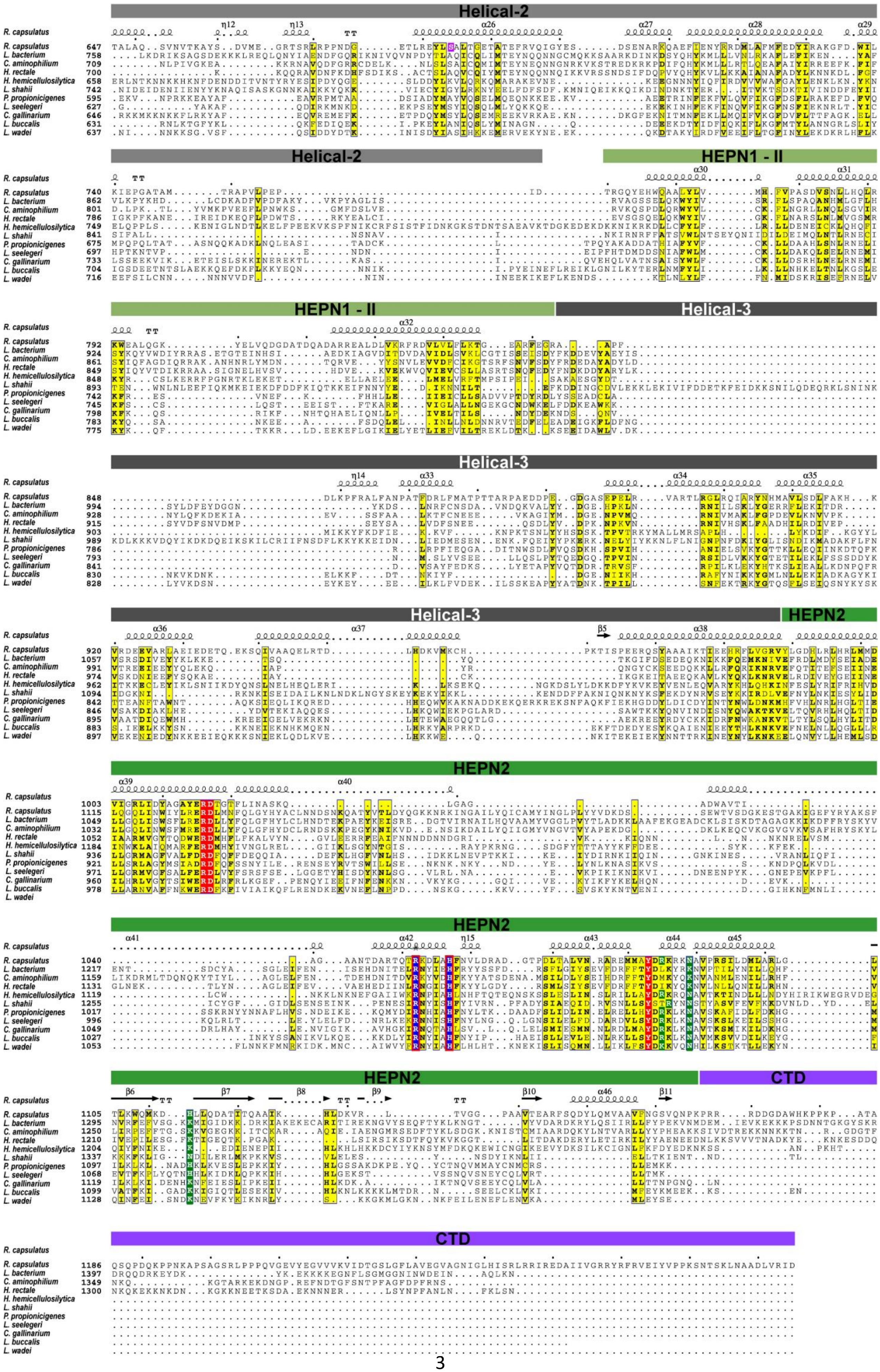
Helical-2
HEPN1 - II
Helical-3
HEPN2
CTD
R. capsulatus
L. bacterium
C. aminophilium
H. rectale
H. hemicellulosilytica
L. shahii
P. propionicigenes
L. seelegeri
C. gallinarium
L. buccalis
L. wadei

**Supporting Figure S2:** Sequence alignment of Cas13a homologs. Secondary structure assignments according to the RcCas13a-crRNA binary complex structure presented in this work. Catalytic residues of the HEPN domains and residues responsible for pre-crRNA processing are highlighted in blue and green, respectively. Amino acids forming sequence-specific contacts with the crRNA are shown in pink. Conserved aromatic and aliphatic regions are shaded yellow, charged residues in red. NCBI RefSeq / Uniprot ID: *R. capsulatus*: WP_013067728.1 / D5AUW0, *L. bacterium*: WP_022785443.1 / P0DPB7, *C. aminophilum*: WP_031473346.1, *E. rectale*: WP_055061018.1, *H. hemicellulosilytica*: WP_103203632.1 / A0A0H5SJ89, *L. shahii*: WP_018451595.1 / P0DOC6, *P. propionicigenes*: WP_013443710.1 / E4T0I2, *L. seeligeri*: WP_012985477.1 / P0DPB8, *C. gallinarum*: WP_034560163.1, *L. buccalis*: WP_015770004.1 / C7NBY4, *L. wadei*: WP_021746003.1 / U2PSH1. Alignment was performed with ClustalX (3) and annotated manually and using ESPript (4).

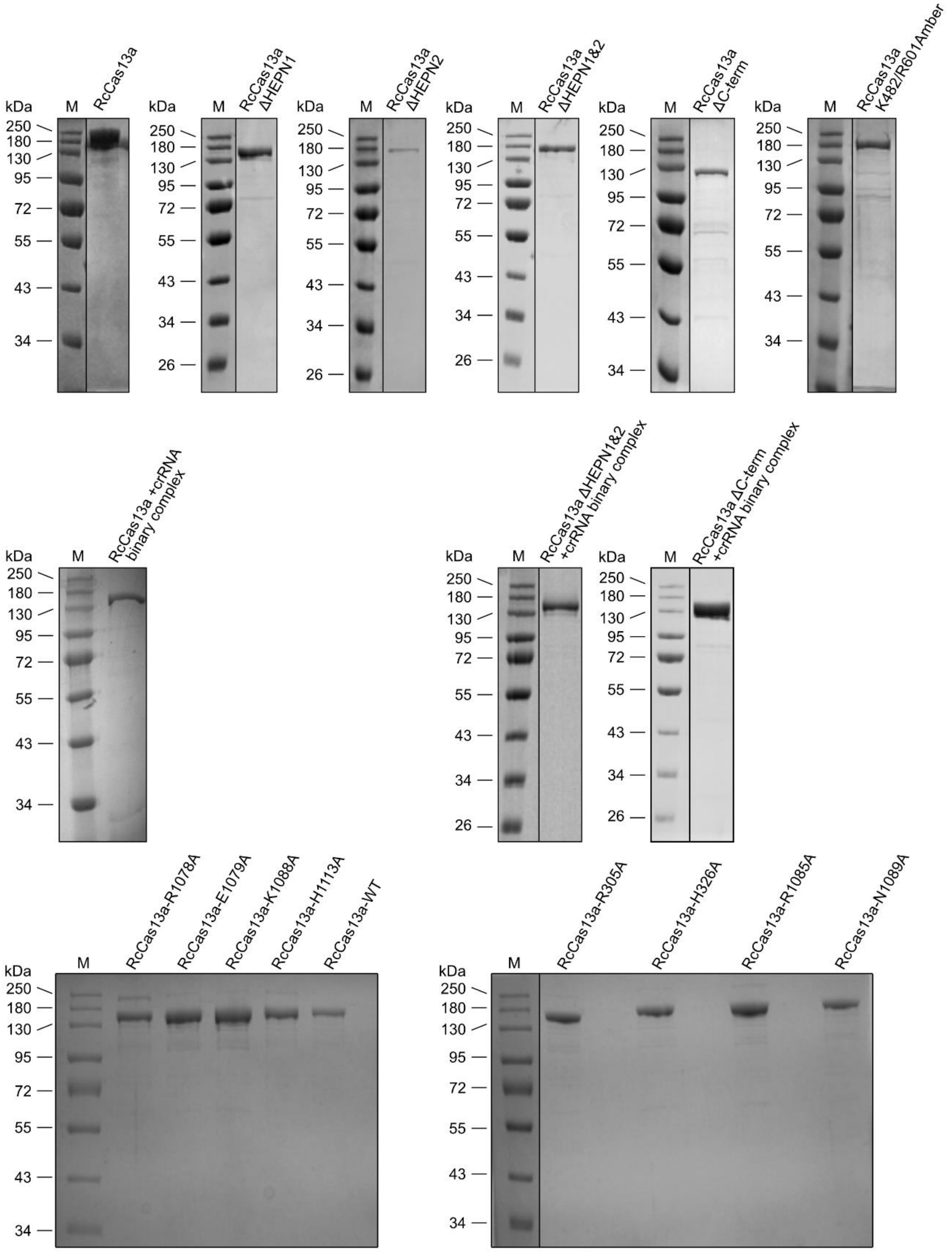


**Supporting Figure S3.** SDS-PAGE analysis (Comassie blue staining) of purified wild type RcCas13a, RcCas13a mutants and the RcCas13a-crRNA complex.

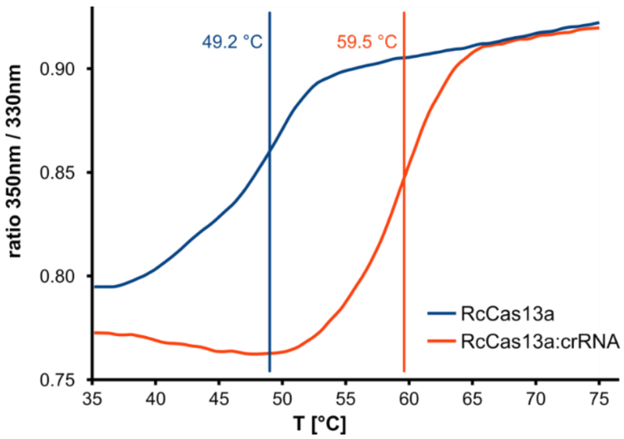


**Supporting Figure S4:** Thermal stability of RcCas13a and the RcCas13a binary complex. Melting curves of apo-RcCas13a (blue) and RcCas13a-crRNA binary complex (orange) determined by nanoDSF (Nanotemper Tycho NT.6) using standard settings.

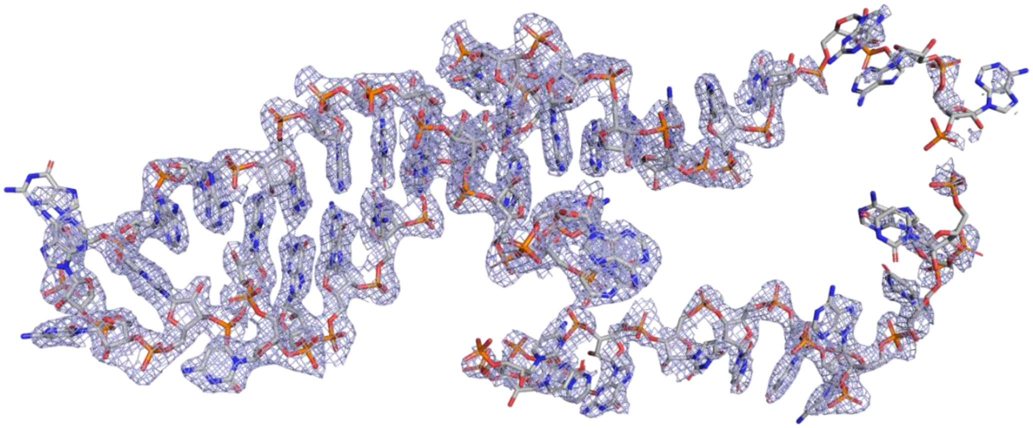

**Supporting Figure S5:** Simulated annealing omit Fo-*D*Fc map of the crRNA chain, contoured at 2.5σ calculated with PHENIX. Parts of the spacer region are flexible and not well defined in the electron density.

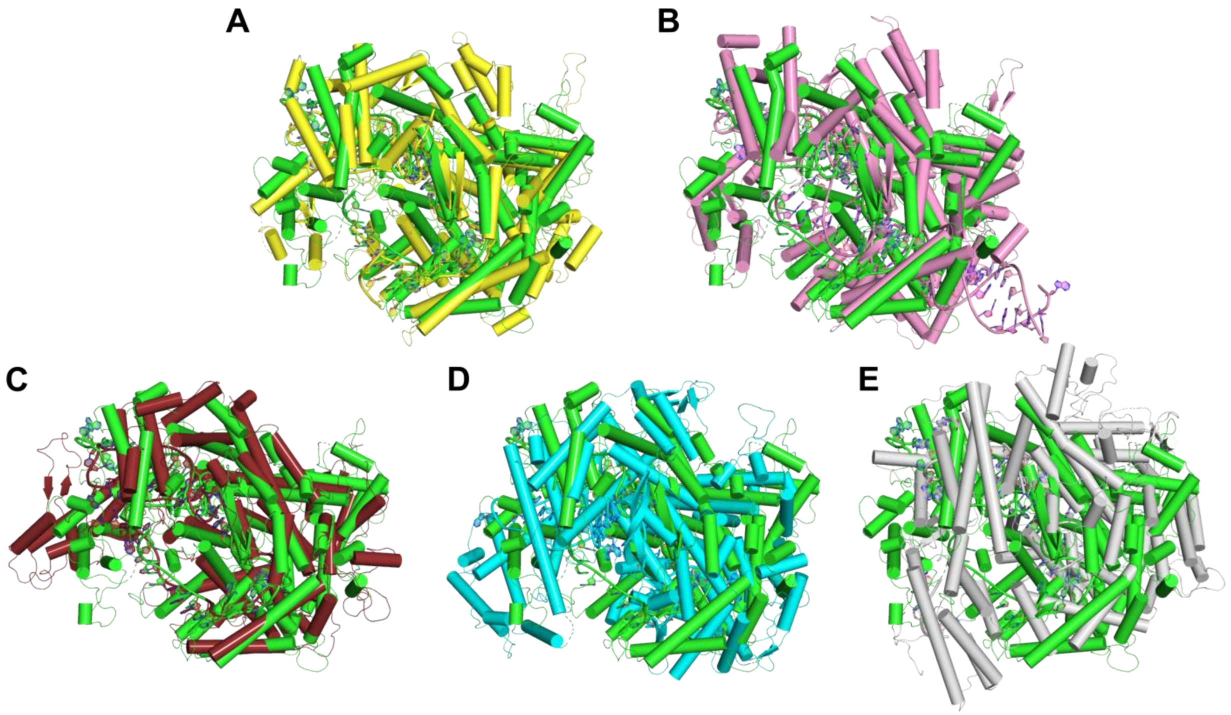


**Supporting Figure S6**: Superposition of RcCas13a with homologous structures. RcCas13a-crRNA binary complex (green) with A) LbuCas13a-crRNA binary complex (yellow) B) LbuCas13a-crRNA-targetRNA ternary complex (pink), C) LseCas13a-crRNA binary complex (brown), D) LbaCas13a-crRNA binary complex (cyan), E) LshCas13a-crRNA binary complex (gray).

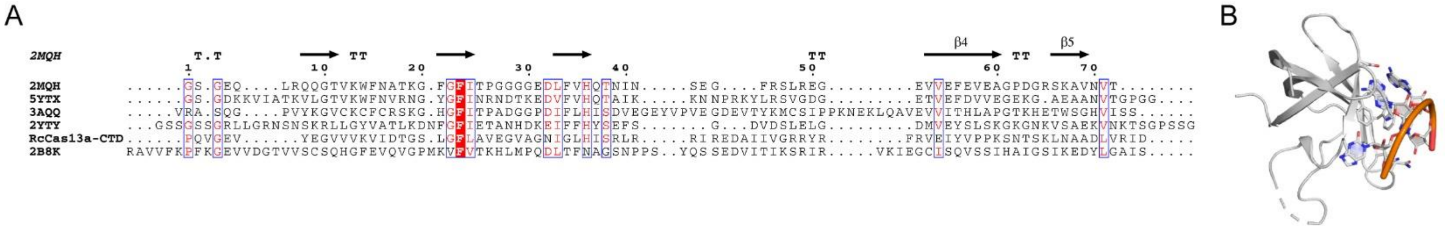


**Supporting Figure S7.** A) Sequence alignment of the RcCas13a-CTD and homologs identified with HHPRED, (5) annotated using ESPript (4). B) Structure of the human Y-box binding protein 1 (YB-1), a member of the cold shock domain (CSD) protein family, in complex with RNA (PDB code 5YTX). RcCas13a might adopt an overall similar folding topology.

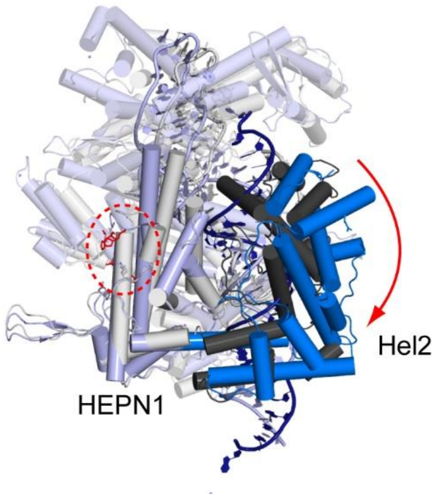


**Supporting Figure S8.** Superposition of the binary (PDB code 5XWY, grey) and ternary complex structures (PDB code 5XWP, blue) of LbuCas13a. Upon target RNA (dark blue) binding the Hel 2 domain rotates about 25° relative to the HEPN1 domain, resulting in the rearrangement of the catalytic residues in the active site (red circle).

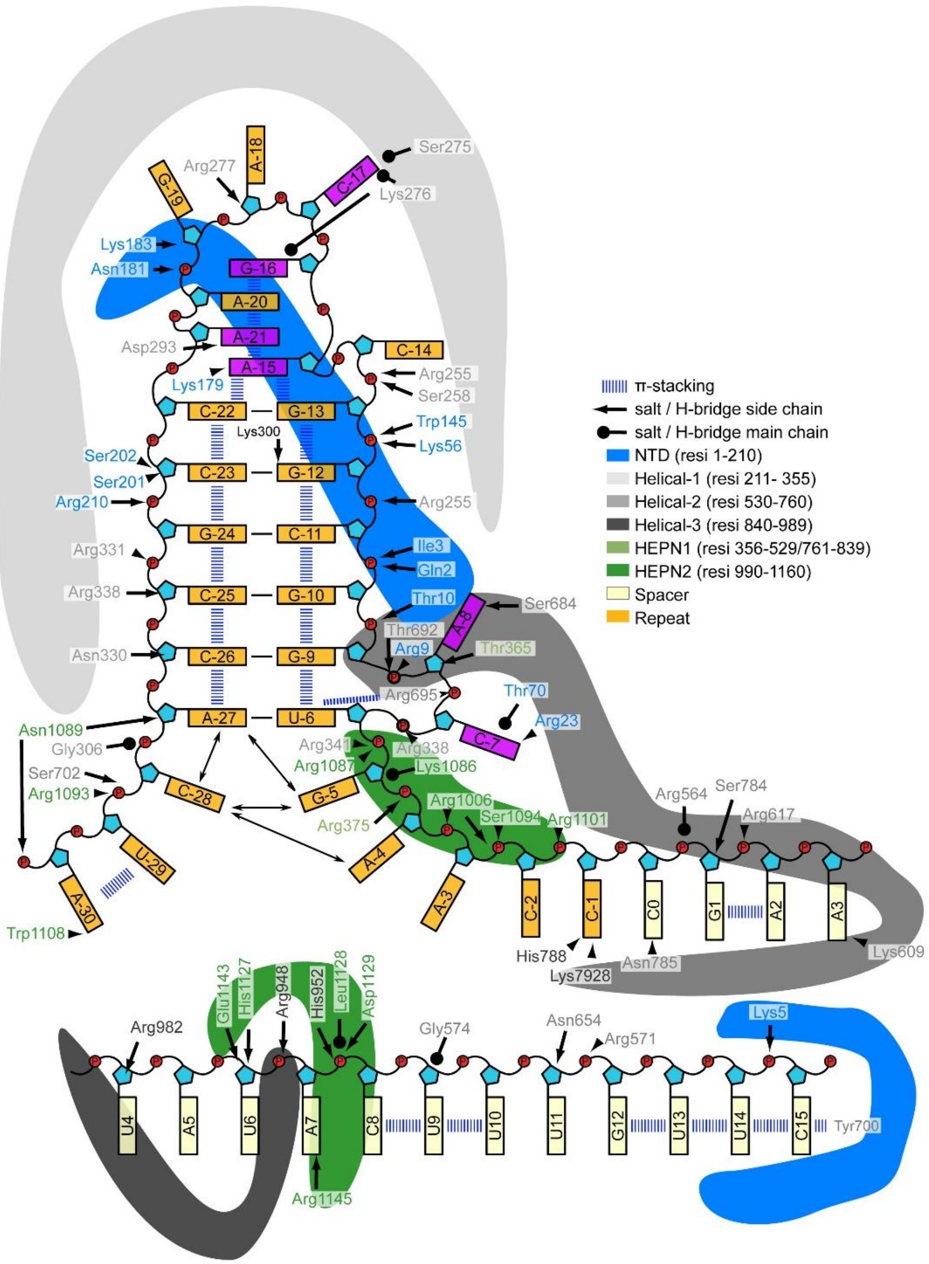

**Supporting Figure S9:** Scheme showing the interactions between crRNA and RcCas13a. Residues from different protein domains are color coded. Base-specific contacts only occur to the stem-loop and 3’ flank. Only 16 nt’s of the spacer are defined in the electron density.

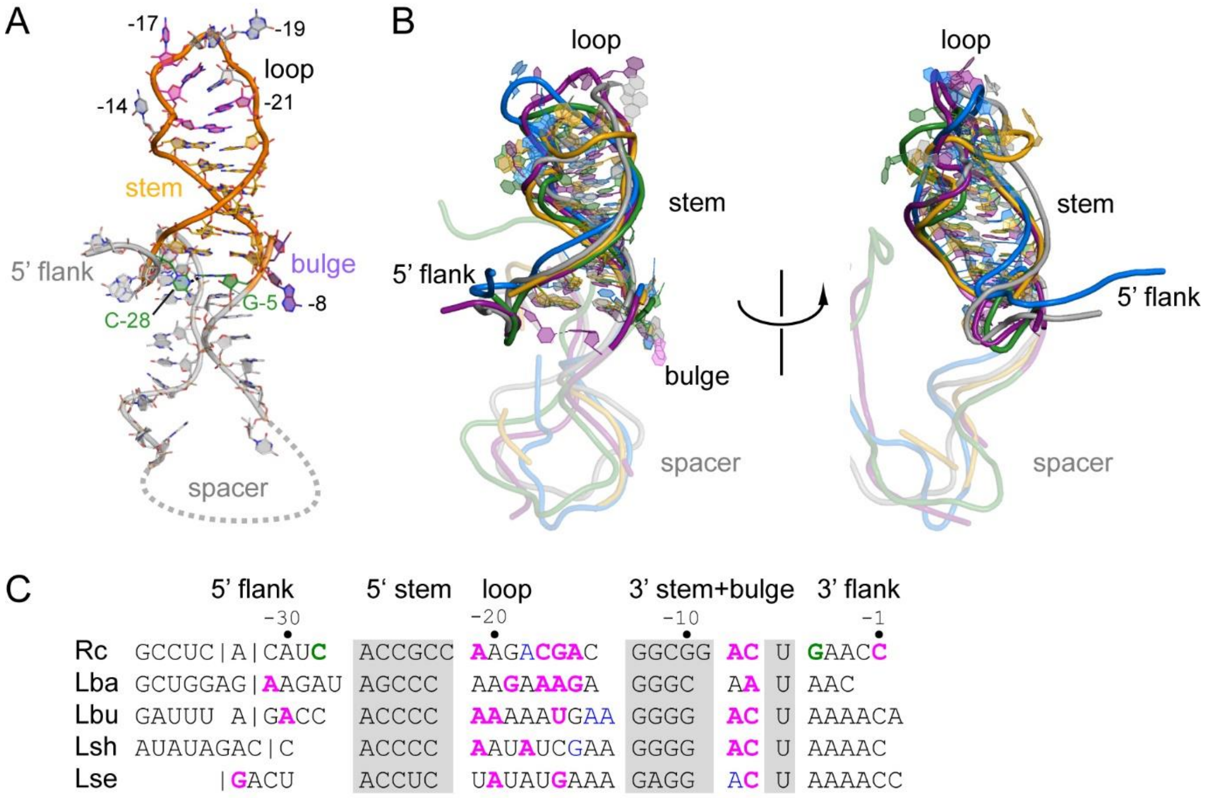


**Supporting Figure S10:** Comparison of crRNAs of the Cas13a-family of endonucleases. A) Structure of the crRNA in the RcCas13a binary complex. The stem of the crRNA is depicted in orange, bulge and residues at the loop (grey) making base specific interactions are highlighted in violet and pink, respectively. The twisted base-pair at the bottom of the stem-loop unique to the RcCas13a bound crRNA is highlighted in green. The spacer and flanking regions are shaded in transparent grey. Disordered residues of the spacer sequence in the X-ray structure of the binary complex are indicated as dashed line. B) Superposition of crRNAs of homologous Cas13a binary complex structures. crRNAs from RcCas13a (purple, this work), LbuCas13a (grey, r.m.s.d. 1.8Å, PDB code 5XWY), LseCas13a (green, r.m.s.d. 1.2Å, PDB code 6VRC)), LbaCas13a (blue, r.m.s.d. 6.2Å, PDB code 5WLH), LshCas13a (golden, r.m.s.d 2.7Å, PDB code 5WTK). The phosphate backbone of the spacer is shown transparent. C) Sequence alignment of crRNA sequences. Shaded grey: bases forming the stem-loop; pink: bases with sequence specific contacts; blue: bases with stacking interactions with the protein; green: twisted base pair unique to the RcCas13a-bound crRNA. | indicates pre-crRNA cleavage position at the 5’ flank.

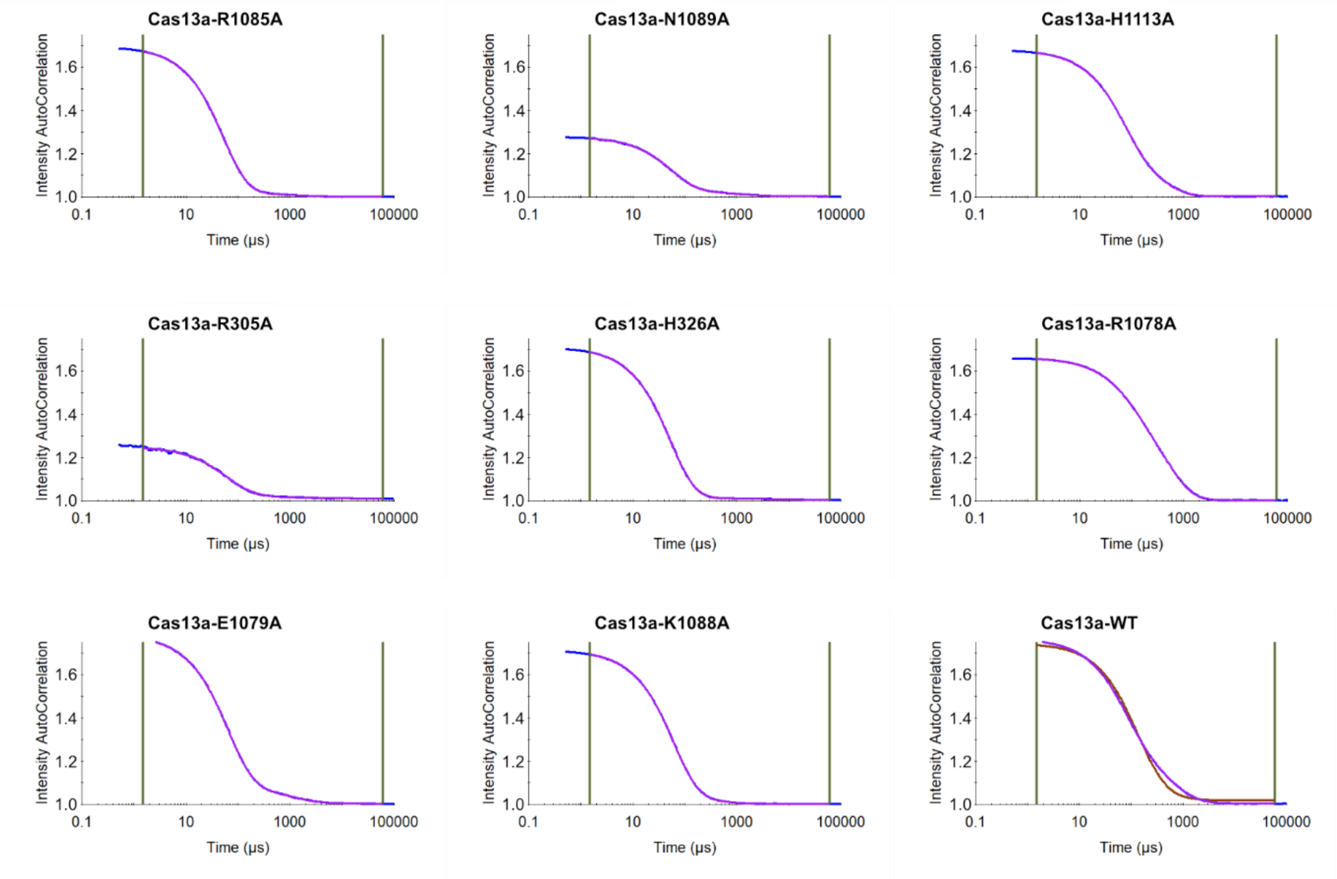


**Supporting Figure S11.** Quality control of purified RcCas13a proteins by DLS-analysis, showing the integrity of all RcCas13a proteins. The observed intensity autocorrelation for the RcCas13a R305A and N1089A mutants are lower compared to the other RcCas13a proteins due to lower protein concentration.

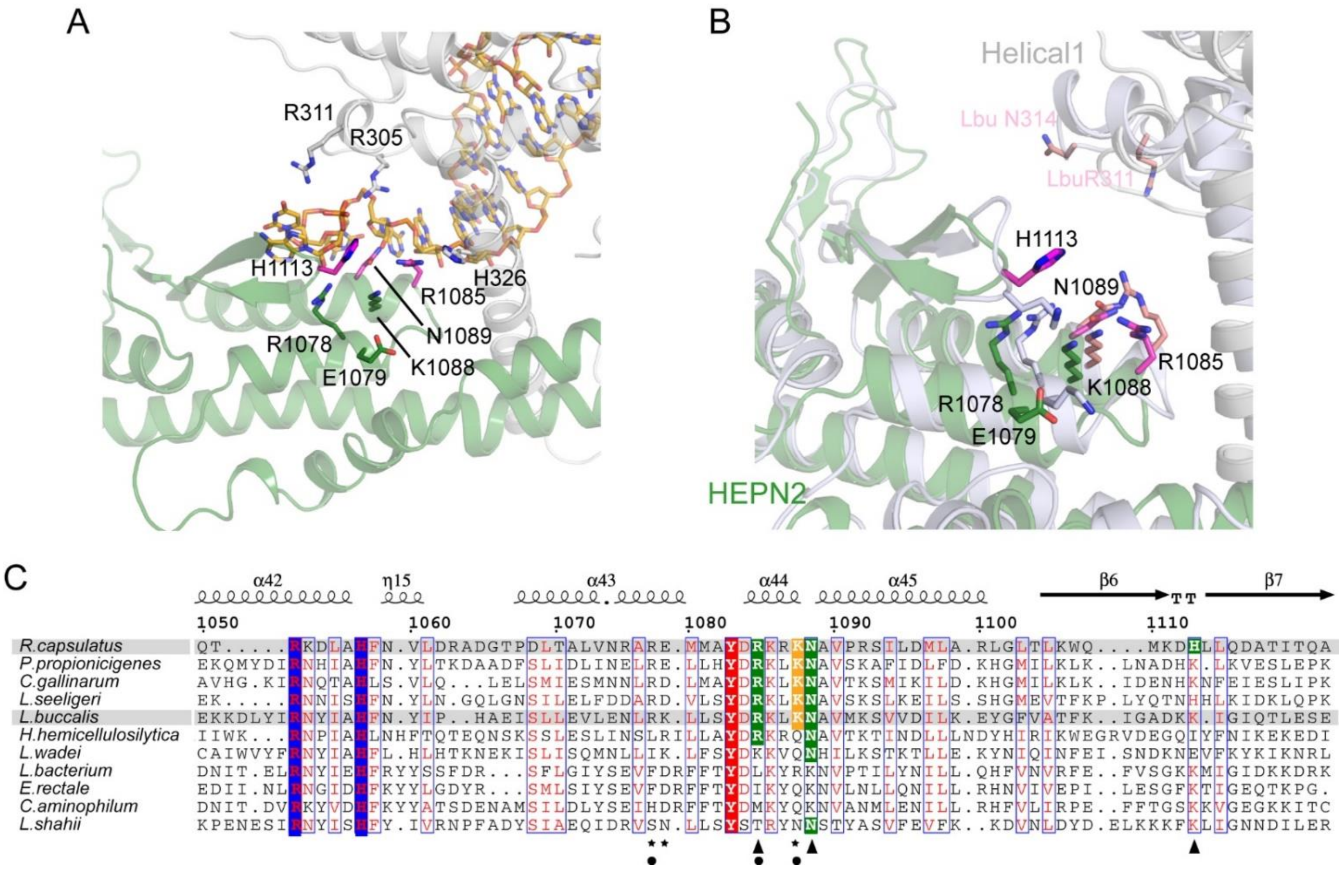


**Supporting Figure S12.** Comparison of RcCas13a and LbuCas13a. A) Close-up view of the crRNA processing site in RcCas13a. Residues that were mutated to Ala are shown as stick model, with the residues important for catalysis highlighted in pink. (HEPN2= green, Helical 1 = grey). B) Superposition of the binary complexes of RcCas13a and LbuCas13a (PDB code 5XWY, light blue) using the main chain atoms of the HEPN2 domain as reference frame. Residues in RcCas13a responsible for pre-crRNA processing are highlighted in pink, the corresponding residues in LbuCas13a are shown in grey. The 5' end of the crRNA in the RcCas13a binary complex is shown as orange stick model. (crRNA in the LbuCas13a binary complex is not shown for clarity). In LbuCas13a two residues (Arg 311, Asn 314) that do not make any contact with the crRNA were identified as crucial for the processing activity. (6) C) Section of a part of the sequence alignment of the HEPN2 domains, with the residues in RcCas13a involved in pre-crRNA processing highlighted in green and marked with an arrow. Residues that where mutated in RcCas13a but did not impact on the processing activity are marked with a star. The residues in LbuCas13a responsible for the pre-crRNA processing (6) are marked with a dot. Arg 1052 and His 1057 involved in target RNA cleavage are highlighted in blue.

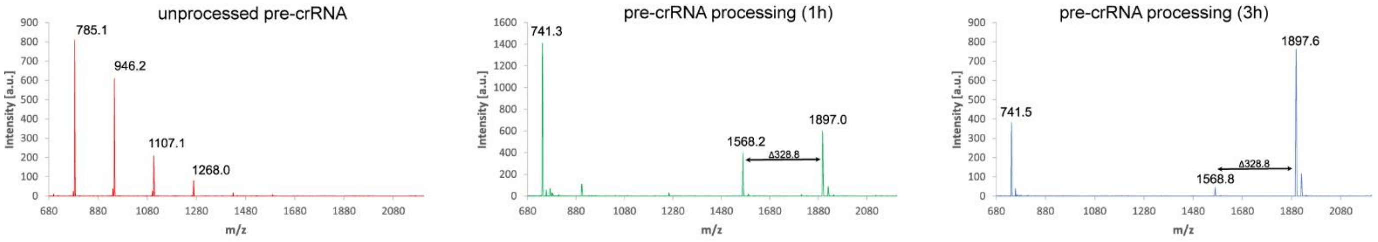


| | 3′-OH | 3′-P | 2′,3′-cyclic-P |
|---|---|---|---|
| 3nt (GCC) | 893.6 | 979.5 | 962.5 |
| 4nt (GCCU) | 1199.7 | 1279.7 | 1262.7 |
| 5nt (GCCUC) | 1504.9 | 1584.9 | 1567.9 |
| 6nt (GCCUCA) | 1834.1 | 1914.1 | 1897.1 |
| 7nt (GCCUCAC) | 2139.3 | 2219.3 | 2202.3 |

*Mass correspond to the average mass
Adenosinmonophosphat 347 Da (-$H_2O$) = 329 Da

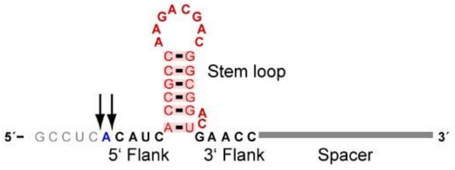


**Supporting Fig. S13.** MALDI-TOF analysis of the pre-crRNA processing reaction products. Spectra of pre-crRNA without incubation with Cas13a (left) and following incubation with RcCas13a (middle and right). Scheme of the pre-crRNA with the cleavage sites indicated by arrows. Table with average masses of possible reaction products calculated with Mongo Oligo Mass Calculator v2.06 (http://rna.rega.kuleuven.be/masspec/mongo.htm).

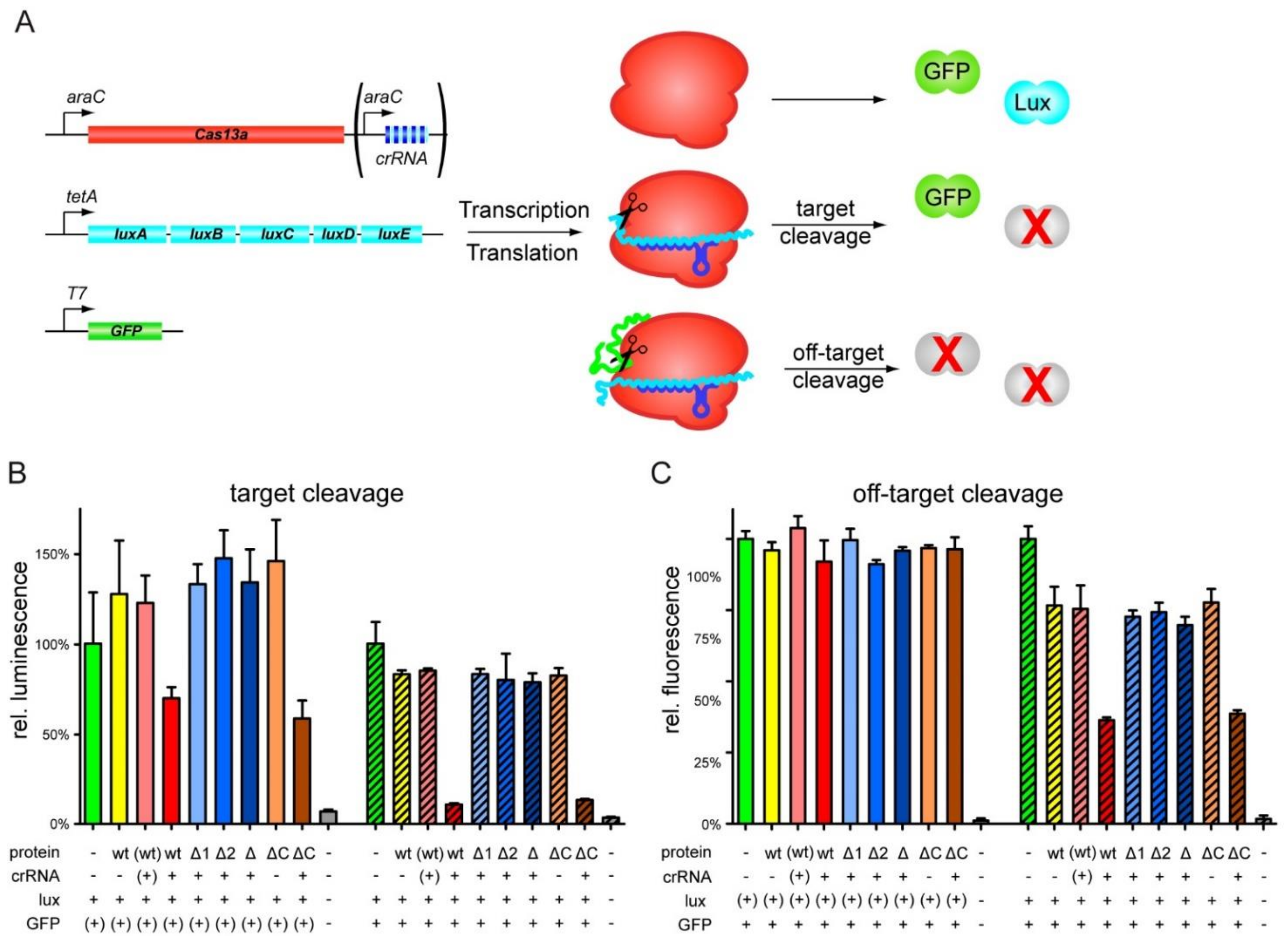


**Supporting Figure S14.** *In vivo* activity of RcCas13a. A) Schematic depiction of the *in-vivo* activity assay. Two reporter gene constructs (*luxABCDE* and *GFP*) as well as expression constructs for RcCas13a variants and crRNA encoded within a CRISPR-locus targeting *luxABCDE* are introduced into *E. coli* cells. Upon expression of RcCas13a and crRNA, luminescence is inhibited by target RNA-cleavage activity of RcCas13a, while fluorescence is prevented by non-sequence specific off-target RNA-cleavage activity. B) Sequence specific target cleavage activity by RcCas13a can be followed by bioluminescence upon luciferase expression reduction. Wt RcCas13a and RcCas13aΔCTD reduce bioluminescence in a crRNA dependent manner (red and brown bars), while all HEPN-mutants lost their impact on luciferase expression (blue bars). This effect is not affected by induction of GFP expression (right, hatched bars). C) Non-sequence specific RNA cleavage was analyzed by measuring GFP-fluorescence upon induction of crRNA and target RNA expression. Wt RcCas13a and RcCas13aΔCTD reduce GFP-fluorescence only upon expression of crRNA and target sequence (red and brown bars). Mutation of any of the catalytic residues in the HEPN-domains leads to an inactive RcCas13a. (blue hatched bars). No impact on GFP-fluorescence can be observed in the absence of *luxABCDE* expression induction (left). RcCas13a and crRNA expression is induced by addition of arabinose to the growth medium. Luciferase and *GFP* expression is controlled by anhydrotetracycline and IPTG, respectively. (+): Encoding gene/expression construct is present in the *E. coli* cells, but was not induced. (-) empty vector control. (Δ1 = Δ HEPN1, Δ2 = Δ HEPN2, Δ = Δ HEPN1+2; ΔC = Δ CTD). Determined values for fluorescence and bioluminescence were corrected by the cell density ($OD_{600}$) and normalized to the empty vector control.

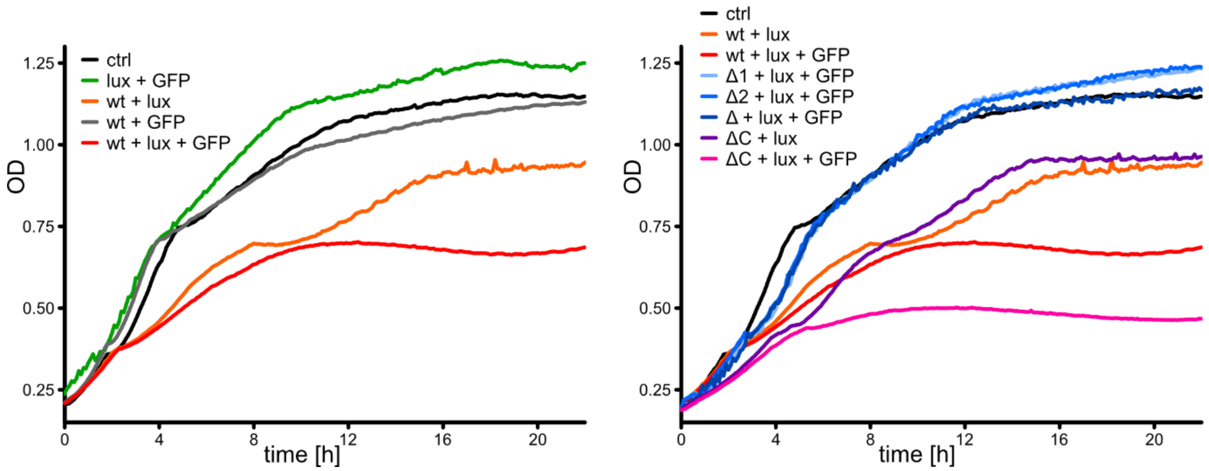


**Supporting Figure S15:** Bacterial growth during *in vivo* activity. Growth curves of RcCas13a and reporter gene expressing *E. coli* JM109 (DE3) over 20 h at 30 °C. ctrl: empty vector control. Induction of expression of active RcCas13a together with crRNA and target RNA leads to a reduction in growth.

**Table S1: Data processing, structure solution and refinement statistics.** Numbers in parentheses correspond to the highest resolution shell.

| | Native PDB code | SeMet |
|---|---|---|
| Wavelength (Å) | 0.966 | 0.979 |
| Resolution range (Å) | 47.1 - 2.2 (2.3 - 2.2) | 47.7 – 2.5 (2.7 - 2.5) |
| Space group | P1 | P1 |
| Unit cell lengths (Å) | 61.4 91.1 136.5 | 62.6 91.2 137.1 |
| Unit cell angles (°) | 90.0 103.7 97.6 | 90.0 103.5 96.7 |
| Total reflections | 504,148 (48,268) | 1,237,991 (69,583) |
| Unique reflections | 141,245 (14,010) | 195,545 (18,034) |
| Multiplicity | 3.6 (3.4) | 6.3 (3.9) |
| Completeness (%) | 97.6 (96.3) | 96.5 (89.1) |
| Mean I/σ(I) | 9.6 (1.2) | 8.9 (1.6) |
| Wilson B-Factor | 41.8 | 411.6 |
| R-merge | 0.085 (0.913) | 0.153 (0.742) |
| R-meas | 0.101 (1.09) | 0.166 (0.861) |
| R-pim | 0.053 (0.585) | 0.064 (0.436) |
| $CC_{1/2}$ | 0.997 (0.726) | 0.995 (0.745) |
| CC* | 0.999 (0.917) | 0.999 (0.924) |
| Reflections used | | |
| in Refinement | 134,180 | |
| for R-free | 7,063 | |
| R-work | 0.218 | |
| R-free | 0.258 | |
| CC(work) | 0.956 (0.802) | |
| CC(free) | 0.950 (0.732) | |
| Number of | | |
| Non-H atoms | 20,527 | |
| macromolecules | 19,714 | |
| ligands | 56 | |
| solvent | 757 | |
| Protein residues | 2,254 | |
| RMS (bonds) | 0.005 | |
| RMS (angles) | 0.88 | |
| Ramachandran | | |
| favored (%) | 98.2 | |
| allowed (%) | 1.75 | |
| outliers (%) | 0.04 | |
| Rotamer outliers | 0.39 | |
| Clashscore | 4.69 | |
| Average B-factor | 56.3 | |
| for macromolecules | 56.5 | |
| for ligands | 71.9 | |
| for solvents | 50.1 | |
| Number of TLS groups | 18 | |

**Table S2: Protein sequence of RcCas13a.** The HEPN active site residues are highlighted in red, residues responsible for pre-crRNA processing in green, and the C-terminal domain (CTD) is highlighted in yellow.

| | |
|---|---|
| RcCas13a | MQIGKVQGRTISEFGDPAGGLKRKISTDGKNRKELPAHLSSDPKALIGQWISGIDKIYRKPDSRK<br>SDGKAIHSPTPSKMQFDARDDLGEAFWKLVSEAGLAQDSDYDQFKRRLHPYGDKFQPADSGAKLK<br>FEADPPEPQAFHGRWYGAMSKRGNDAKELAAALYEHLHVDEKRIDGQPKRNPKTDKFAPGLVVAR<br>ALGIESSVLPRGMARLARNWGEEEIQTYFVVDVAASVKEVAKAAVSAAQAFDPPRQVSGRSLSPK<br>VGFALAEHLERVTGSKRCSFDPAAGPSVLALHDEVKKTYKRLCARGKNAARAFPADKTELLALMR<br>HTHENRVRNQMVRMGRVSEYRGQQAGDLAQSHYWTSAGQTEIKESEIFVRLWVGAFALAGRSMKA<br>WIDPMGKIVNTEKNDRDLTAAVNIRQVISNKEMVAEAMARRGIYFGETPELDRLGAEGNEGFVFA<br>LLRYLRGCRNQTFHLGARAGFLKEIRKELEKTRWGKAKEAEHVVLTDKTVAAIRAIIDNDAKALG<br>ARLLADLSGAFVAHYASKEHFSTLYSEIVKAVKDAPEVSSGLPRLKLLLKRADGVRGYVHGLRDT<br>RKHAFATKLPPPPAPRELDDPATKARYIALLRLYDGPFRAYASGITGTALAGPAARAKEAATALA<br>QSVNVTKAYSDVMEGRTSRLRPPNDGETLREYLSALTGETATEFRVQIGYESDSENARKQAEFIE<br>NYRRDMLAFMFEDYIRAKGFDWILKIEPGATAMTRAPVLPEPIDTRGQYEHWQAALYLVMHFVPA<br>SDVSNLLHQLRKWEALQGKYELVQDGDATDQADARREALDLVKRFRDVLVLFLKTGEARFEGRAA<br>PFDLKPFRALFANPATFDRLFMATPTTARPAEDDPEGDGASEPELRVARTLRGLRQIARYNHMAV<br>LSDLFAKHKVRDEEVARLAEIEDETQEKSQIVAAQELRTDLHDKVMKCHPKTISPEERQSYAAAI<br>KTIEEHRFLVGRVYLGDHLRLHRLMMDVIGRLIDYAGAYERDTGTFLINASKQLGAGADWAVTIA<br>GAANTDARTQTRKDLAHFNVLDRADGTPDLTALVNRAREMMAYDRKRKNAVPRSILDMLARLGLT<br>LKWQMKDHLLQDATITQAAIKHLDKVRLTVGGPAAVTEARFSQDYLQMVAAVFNGSVQNPKPRRR<br>DDGDAWHKPPKPATAQSQPDQKPPNKAPSAGSRLPPPQVGEVYEGVVVKVIDTGSLGFLAVEGVA<br>GNIGLHISRLRRIREDAIIVGRRYRFRVEIYVPPKSNTSKLNAADLVRID |

**Table S3: RNA sequences.** FAM= 6-carboxyfluorescein

| Name | Sequence (5' to 3' directionality) |
|---|---|
| pre-crRNA for *in vitro* processing | GCCUCACAUCACCGCCAAGACGACGGCGGACUGAACCGGGUGUCGC<br>CCUCGAACUUCACCUCGGC-3′-FAM |
| crRNA for crystallization | GCCUCACAUCACCGCCAAGACGACGGCGGACUGAACCUUCAUUACC<br>UCUGUUUG |
| CRISPR locus for *in vivo* activity assay | GACCUGAGCCUCACAUCACCGCCAAGACGACGGCGGACUGAACCUU<br>CAUUACCUCUGUUUGAGAAAAUUGGGGCCUCACGUCACCGCCAAGA<br>CGACGGCGGACUGAACCCGAACUAUACUUUGUUCUUGAACAGUUGG<br>CAUCACAUCACCGCCAAGACGACGGCGGACUGAACCCACCAAAAUU<br>AAUGGAUUGCACUAAAUCGCCUCACAUCACCGCCAAGACGACGGCG<br>GACUGAACCGAUCCAUCCUGCGGGCAAAACCAACGCGCCUCACAUC<br>ACCGCCAAGAUGACGGCGGGAACCCAACCACACUAAUGGAUCGCUC<br>GAAAAAAUCUGCAAAC |

**Table S4: Plasmids used in this study.***RcCas13a single mutants: R305A, R326A, R1078A, E1079A, K1088A, R1085A, N1089A, H1113A

| Vector name | Application | Selection marker | Source / Construction |
|---|---|---|---|
| pET24-RcCas13a + crRNA | Protein expression (crystallisation) | Kanamycin | This work |
| p2CT-His-MBP-TEV-RcCas13a | Protein expression | Ampicillin | Addgene #91870 |
| p2CT-His-MBP-TEV-RcCas13a* | Protein expression | Ampicillin | This work<br>*Wt and RcCas13a single mutants: R305A, R326A, R1078A, E1079A, K1088A, R1085A, N1089A, H1113A |

| | | | |
|---|---|---|---|
| p2CT-RcCas13a ΔHEPN1 | Protein expression | Ampicillin | This work p2CT-RcCas13a R464A/H469A |
| p2CT-RcCas13a ΔHEPN2 | Protein expression | Ampicillin | This work p2CT-RcCas13a R1052A/H1057A |
| p2CT-RcCas13a ΔHEPN1+2 | Protein expression | Ampicillin | This work p2CT-RcCas13a R464A/H469A R1052A/H1057A |
| pASK-IBA5-luxABCDE | *In vivo* activity assay | Ampicillin | This work |
| pACYC-eGFP-ssrA | *In vivo* activity assay | Chlor-amphenicol | This work |
| pBAD-RSF1031K-RcCas13a | *In vivo* activity assay | Kanamycin | This work |
| pBAD-RSF1031K-RcCas13a + crRNA | *In vivo* activity assay | Kanamycin | This work |
| pBAD-RSF1031K-RcCas13a ΔHEPN1 | *In vivo* activity assay | Kanamycin | This work pBAD-RSF1031K-RcCas13a R464A/H469A |
| pBAD-RSF1031K-RcCas13a ΔHEPN2 | *In vivo* activity assay | Kanamycin | This work pBAD-RSF1031K-RcCas13a R1052A/H1057A |
| pBAD-RSF1031K-RcCas13a ΔHEPN1+2 | *In vivo* activity assay | Kanamycin | This work pBAD-RSF1031K-RcCas13a R464A/H469A/ R1052A/H1057A |
| pBAD-RSF1031K-RcCas13a ΔHEPN1 + crRNA | *In vivo* activity assay | Kanamycin | This work pBAD-RSF1031K-RcCas13a + crRNA R464A H469A |
| pBAD-RSF1031K-RcCas13a ΔHEPN2 + crRNA | *In vivo* activity assay | Kanamycin | This work pBAD-RSF1031K-RcCas13a + crRNA R1052A H1057A |
| pBAD-RSF1031K-RcCas13a ΔHEPN1+2 + crRNA | *In vivo* activity assay | Kanamycin | This work pBAD-RSF1031K-RcCas13a + crRNA R464A/H469A R1052A/H1057A |

**Table S5: List of primers used in this study.**

| Internal number | Name | Sequence (5' to 3' directionality) |
|---|---|---|
| 607 | LK-22_CRISPR-loci-SalI_fw | gatcgtcgacgatccaatcgag |
| 608 | LK-23_CRISPR-loci-SbfI_rev | gatccctgcaggtcgaaaaagg |
| 609 | LK-24_C2c2-seq1_fw | tttgccgctcctcg |
| 610 | LK-25_C2c2-seq2_fw | ttccgcctgtttgc |
| 611 | LK-26_C2c2-seq3_rev | aaccaaagcggaacc |
| 631 | RcDoudna | taatacgactcactatagtcacatcaccgccaagacgacggcggac tgaaccgg cttaccccatacccaagaaaggaacatcatg |
| 632 | CRISPRrev | catgatgttcctttcttgggtatg |
| 641 | LK-20_C2c2-XbaI_fw_2 | gatctctagacatatgcagattggcaagg |
| 642 | LK-21-C2c2-BsaI/XhoI_rev_2 | gatcctcgaggagacctcagtcgatccgcaccag |

| | | |
|---|---|---|
| 648 | C2c2 3' seq fw | gtaacacgtcgaagc |
| 649 | pLIKE loci rc seq | cacgaaacaataattgg |
| 653 | LK-27_C2c2-GA_fw | caacaccaattagttaaggaggattctagacatatgcagattggca aggttc |
| 654 | LK-28_C2c2-GA_rev | ttaggcgggctgcccggggacgtcctcgaggagacctcagtcgatc cgcaccagatc |
| 655 | LK-29_loci-GA_fw | gccccttttttcatgaaactagttcgtcgacgatccaatcgagtc |
| 656 | LK-30_loci-GA_rev | cggaattagcttgcatgcctgcaggtcgaaaaaggggcagccc |
| 665 | LK-31_NcoI-blaI_fw | gatcccatggatgaaaaaaatacctcaaatctc |
| 666 | LK-32_blaI-HindIII_rev | gatcaagctttcattccttctttctgttc |
| 667 | LK-33_C2c2-TEV-correction_fw | cgagtgcggccgcaaggctctggaaatacaagttttcg |
| 668 | LK-34_C2c2-TEV-correction_rev | cgaaaacttgtatttccagagccttgcggccgcactcg |
| 669 | LK-35_C2c2-pBAD_fw | tttaagaaggagatatacatacccatggatgcagattggcaaggtt c |
| 670 | LK-36_C2c2-pBAD_rev | ctgaggtacctcagtggtggtggtggtg |
| 671 | LK-37_C2c2-ara/loci_rev | tagggataggcttaccttcaagctttcagtggtggtggtggtg |
| 672 | LK-38_ara/loci-C2c2_fw | ccaccactgacggtacctcagagaagaaac |
| 673 | LK-39_ara/loci-pBAD_rev | tagggataggcttaccttcaagctttctagagtttgcagattttt c |
| 674 | LK-40_lux-pACYC_fw | accatcatcaccacagccaggatccatgaaatttggaaactttttg c |
| 675 | LK-41_lux-pACYC_rev | ctgcaggcgcgccgagctcgaattctcaactatcaaacgcttcg |
| 681 | LK-42_pBAD-rev-seq | ctctcatccgccaaaac |
| 682 | LK-43_C2c2-5'-seq-rev | gatcgcttgacagatgag |
| 693 | ACYCDuetUP1 | ggatctcgacgctctccct |
| 694 | DuetDown1 | gattatgcggccgtgtacaa |
| 695 | LK-44_pBAD-C2c2_fw2 | tttaagaaggagatatacatacccatggatgcagattggcaaggtt caaggg |
| 696 | LK-45_C2c2-pBAD_rev2 | tagggataggcttaccttcaagctttcagtgtgtgtggtggtggtg |
| 697 | LK-46_C2c2-ara/loci_rev2 | ggtttcttctctgaggtaccgtcagtggtggtggtggtggtg |
| 698 | LK-47_ara/loci-C2c2_fw2 | caccaccaccaccaccactgacggtacctcagagaagaaac |
| 704 | LK-48_lux-pACYC_rev2 | cagcggtttctttaccagactcgagtcaactatcaaacgcttcg |
| 703 | LK-45_C2c2-pBAD_rev3 | tagggataggcttaccttcaagctttcagtggtggtggtggtggtg c |
| 705 | LK-49_pBADRSF1031K_rev | ctctagaggatccccggg |
| 706 | LK-50_pBADRSF1031K_fwd | aagcttggctgttttggc |
| 707 | LK-51_C2c2_into_pBADRSF1031K_fwd | tacccggggatcctctagagatgcagattggcaaggttc |
| 708 | LK-52_C2c2_into_pBADRSF1031K_rev | ccgccaaaacagccaagcttgcttcctttcgggctttg |
| 709 | LK-53_C2c2-loci_into_pBADRSF1031K_rev | tgaggtaccggcttcctttcgggctttg |
| 710 | LK-54_ara/loci-lux_into_pBADRSF1031K_fwd | gaaaggaagccggtacctcagagaagaaac |
| 711 | LK-55_ara/loci-lux_into_pBADRSF1031K_rev | ccgccaaaacagccaagctttctagagtttgcagatttttc |
| 712 | LK-56_pASK-rev | ccgagctcgaattcgggac |
| 713 | LK-57_pASK-fwd | cggggatccctcgaggtc |
| 714 | LK-58_luxA-E_into_pASK_fwd | ggtcccgaattcgagctcggatgaaatttggaaactttttgc |
| 715 | LK-59_luxA-E_into_pASK_rev | tcgacctcgagggatccccgtcaactatcaaacgcttcg |
| 716 | LK-60_BamHI_luxA_fwd | gatcggatccatgaaatttggaaactttttgc |
| 717 | LK-61_EcoRI_luxE_rev | gatcagaattctcaactatcaaacgcttcg |
| 718 | pASK seq rev | cgcagtagcggtaaacg |
| 719 | pBAD RSF1031 seq rev | ggtcaggtgggaccacc |

| | | |
|---|---|---|
| 730 | LK-62_HindIII-loci | gatcaagcttcggtacctcagagaagaaac |
| 731 | LK-63_loci-XmnI | gatcgaagatcttctctagagtttgcagatttttc |
| 734 | C2c2 3' colony PCR | gccagatcaaaaaccgcccaac |
| 753 | LK-64_QC-HEPN1_fw | ctgcgcggttgcgcgaaccagacctttgcccttggtgcc |
| 754 | LK-65_QC-HEPN1_rev | gcaccaagggcaaaggtctggttcgcgcaaccgcgcag |
| 755 | LK-66_QC-HEPN2_fw | cgcgcgcacccaaaccgccaaagaccttgcggctttcaatgtgc |
| 756 | LK-67_QC-HEPN2_rev | gcacattgaaagccgcaaggtctttggcggtttgggtgcgcgcgtc |
| 764 | LK-72_p2CT-C2c2-seq1 | attttatcgatcccactgatcc |
| 765 | LK-73_p2CT-C2c2-seq2 | attcaaacttatttcgtcgtcg |
| 766 | LK-74_p2CT-C2c2-seq3 | aaggcaaaagaggcgg |
| 767 | LK-75_p2CT-C2c2-seq4 | cctgaacctattgacacacg |
| 768 | LK-76_p2CT-C2c2-seq5 | acgtttcttattaatgcttcaaagc |
| 769 | MBP-fw | gatgaagccctgaaagacgcgcag |
| 770 | LK-77_rrnB-T2_rev | ccatccgtcaggatggcc |
| 771 | LK-78_pET-NcoI | gatcccatgggagttggctgctgccac |
| 772 | LK-79_pET-T7-lacO-NheI | gatcgctagcggggaattgttatccgctcacaattcccctatagtg agtcgt attaaagcttcctttcgggctttg |
| 773 | LK-80_loci-NcoI | gatcccatggtctagagtttgcagatttttcg |
| 53 | pASK-Iba5for | gagttattttaccactccct |
| 61 | T7-in-vitro-TX | gatgtaatacgactcactatag |
| 888 | LK-119 QC p2CT HEPN1 fw | cgttatttacgtgggtgtgctaaccagaccttcgctctgggcgcac gtgctg |
| 889 | LK-120 QC p2CT HEPN1 rev | cagcacgtgcgcccagagcgaaggtctggttagcacacccacgtaa ataacg |
| 890 | LK-121 QC p2CT HEPN2 fw | gctcgtacacaaacggctaaggaccttgcggcctttaatgtgttgg |
| 891 | LK-122 QC p2CT HEPN2 rev | ccaacacattaaaggccgcaaggtccttagccgtttgtgtacgagc |
| 894 | LK-123 p2CT-C2c2-seq6 | ggattgaccccatggg |
| 917 | LK-134_ITX11-fw | ccaagtaatacgactcactatag |
| 918 | LK-135_ITX12-rev | gccgaggtgaagttc |
| 919 | LK-136_ITX13-crRNA-wt | ccaagtaatacgactcactataggcctcacatcaccgccaagacga cggcggact gaaccgggtgtcgccctcgaacttcacctcggc |
| 895 | PM-1 SLIM fw1 C-term pet24 | cttcaatggcagcgtccaggaaaacttgtatttccagagccttg |
| 896 | PM-2 SLIM rev1 C-term pet24 | acggccgccac |
| 897 | PM-3 SLIM fw2 C-term pet24 | gaaaacttgtatttccagagccttg |
| 898 | PM-4 SLIM rev2 C-term pet24 | ctggacgctgccattg |
| 899 | PM-5 SLIM fw1 C-term p2CT | gctgtatttaatggaagtgtacagtaataacattggaagtggataa c |
| 900 | PM-6 SLIM rev1 C-term p2CT | agctaccatctgcaag |
| 901 | PM-7 SLIM fw2 C-term p2CT | taataacattggaagtggataac |
| 902 | PM-8 SLIM rev2 C-term p2CT | ctgtacacttccattaaatacag |
| 903 | PM-9 SLIM fw3 C-term p2CT | cttccaatccaatgcaatgcaagttggtgaggtgtac |
| 904 | PM-10 SLIM rev3 C-term p2CT | tacaggttttcctcgatcc |
| 905 | PM-11 SLIM fw4 C-term p2CT | caagttggtgaggtgtac |
| 906 | PM-12 SLIM rev4 C-term p2CT | cattgcattggattggaag |
| 926 | PM-13 SLIM fw1 C-term CB453 | cttcaatggcagcgtccagtaactcgagtaaggatctccag |
| 927 | PM-14 | cacctcgacaaggtcagg |
| 928 | PM-15 | gatgaagccctgaaagacg |
| 956 | LK-148 Cas13a-stop-Cterm fw | ggcagcgtccagtaaccaaagccacgc |

| | | |
|---|---|---|
| 957 | LK-149 Cas13a-stop-Cterm rev | cgtggctttggttactggacgctgc |
| 1006 | LK-175_pACYC-fw | ggtaccctcgagtctggtaaag |
| 1007 | LK-176_pACYC-rev | atgtatatctccttcttatacttaactaatatactaagatg |
| 1008 | LK-177_eGFP-fw | tataagaaggagatatacatatggtttc |
| 1009 | LK-178_eGFP-rev | ttaccagactcgagggtac |
| 1010 | LK-179_eGFP-colony_fw | gccgaggtgaagttcg |
| 1032 | LK-187_deGFP_fw | tataagaaggagatatacatatggagcttttcactggcgttg |
| 1033 | LK-188_deGFP_rev | ttaccagactcgagggtaccttagatcccggcggcggtc |
| 1034 | LK-189_deGFP-ssrA_rev | ttaccagactcgagggtaccttaagctgccagagcgtagttttcat cattagcagccggccggatcccggcggcggtc |
| 1105 | QC-fw-Cas13a-R1078A | ccgcgctgcggagatgatggcttac |
| 1106 | QC-rv-Cas13a-R1078A/E1079A | ttcaccaacgcggttaaatcg |
| 1107 | QC-fw-Cas13a-E1079A | ccgcgctcgtgcgatgatggctt |
| 1106 | QC-rv-Cas13a-R1078A/E1079A | ttcaccaacgcggttaaatcg |
| 1108 | QC-fw-Cas13a-K1088A | ccgtaaacgcgcgaatgctgtac |
| 1109 | QC-rv-Cas13a-K1088A | tcgtaagccatcatctcac |
| 1110 | QC-fw-Cas13a-H1113A | aatgaaagatgcgttgttgcaagacgctactattac |
| 1111 | QC-rv-Cas13a-H1113A | tgccattttaaagtcaatc |
| 1133 | QC-fw-Cas13a-R305A | tttgtgcgcggcgggtaagaatg |
| 1134 | QC-rv-Cas13a-R305A | cgcttgtaagttttctttac |
| 1135 | QC-fw-Cas13a-H326A | attgatgcgcgcgacccatgaaaacc |
| 1136 | QC-rv-Cas13a-H326A | gctaacaattccgttttg |
| 1137 | QC-fw-Cas13a-R1085A | ggcttacgacgcgaaacgcaagaatgctgtac |
| 1138 | QC-rv-Cas13a-R1085A | atcatctcacgagcgcgg |
| 1139 | QC-fw-Cas13a-N1089A | taaacgcaaggcggctgtacctcg |
| 1140 | QC-rv-Cas13a-N1089A | cggtcgtaagccatcatc |
